# High sensitivity intrinsic optical signal imaging through flexible, low-cost adaptations of an upright microscope

**DOI:** 10.1101/2022.11.08.515742

**Authors:** Brenda Vasquez, Baruc Campos, Ashley Cao, Aye Theint Theint, William Zeiger

## Abstract

Intrinsic optical signal imaging (IOSI) is a staple technique in modern neuroscience. Pioneered over thirty years ago, IOSI allows macroscopic mapping of neuronal activity throughout the cortex. The technique has been used to study sensory processing and experience-dependent plasticity, and is often used as an adjunctive procedure to localize cortical areas for subsequent targeting by other imaging or physiology techniques. Despite the ubiquity of IOSI in neuroscience there are few commercially available IOSI systems. As a result, investigators have typically resorted to building their own imaging systems. Over the years, simplified systems built on existing microscope platforms have been proposed. Still, these often require additional, sometimes costly, custom-built hardware, which presents challenges for investigators without significant experience in optics or microscopy design. Here we present a straightforward set of adaptations that can be applied to any standard upright microscope, using readily available, inexpensive, commercial parts for illumination, optics, and signal detection, that enables high sensitivity IOSI. Using these adaptations, we are able to readily map sensory-evoked signals across the somatosensory and visual cortex, including single-whisker barrel cortical activity maps in mice. We show that these IOSI maps are highly reproducible across animals and can be used to study plasticity mechanisms in the somatosensory cortex. We also provide open-source applications to control illumination and analyze raw data to generate activity maps. We anticipate these resources will be particularly useful for neuroscience investigators from broad technical backgrounds looking to add IOSI capabilities to an existing microscope on a budget.

## Introduction

Intrinsic optical signal imaging (IOSI) is a macroscopic imaging technique that can indirectly map neuronal activity by measuring changes in intrinsic signals^1^. Depending on the wavelength of light used for illumination, intrinsic signals can arise from a variety of sources, including changes in blood volume and blood flow, alterations in hemoglobin saturation due to oxygen consumption, neurotransmitter release, ion and water movement, and fluctuations in the volume of neuronal cell bodies and capillaries^1–3^. Pioneered in the 1980s^4^, IOSI has since become a staple technique in neuroscience. IOSI has been used to uncover the functional architecture of the cortex, including the retinotopic mapping of the visual cortex^5,6^, somatotopic mapping of the primary somatosensory cortex^7,8^, tonotopic mapping of the auditory cortex^9,10^, and even odorant mapping of the olfactory cortex^11^. It has also been critical for studies of cortical plasticity, such as the plasticity of visual and sensory maps during development^12,13^, in response to sensory deprivation^14–17^ or neurotrophic factors^18^, or after environmental enrichment^19,20^. More recently, IOSI has been used to study the pathogenesis of neuropsychological disorders as diverse as migraines^21^, seizures^22^, stroke^23–25^, and neurodevelopmental disorders^26,27^. From a practical standpoint, IOSI can also be used to localize cortical areas for subsequent targeted interventions like viral vector injections^28^, two-photon imaging^29,30^, in vivo electrophysiology^31^, or focal lesions^32^.

To conduct IOSI, one needs a source of light to illuminate the cortex, optics to collect and focus light reflected from the cortical surface, and a detector to record and quantify the reflected light^1,2^. Over the years, and as technology has advanced, many different approaches to assembling these basic hardware components have been described. Most of these have been custom-built by individual labs as IOSI set-ups have largely not been commercialized. Early set-ups for IOSI utilized a traditional upright microscope housing with arrays of photodetectors^4,33^. This was later largely supplanted by sensitive CCD cameras and a tandem lens macroscope configuration, which improved light collection efficiency by orders of magnitude^34^. Illumination likewise has largely shifted from broadband light sources with specific bandpass emission filter^34,35^ to simpler wavelength-specific light-emitting diodes (LEDs)^28,36^. A few protocols have been published over the years detailing the set-up of IOSI rigs, but these still often require custom machined parts, manually wired circuitry, or expensive commercial hardware such as power supplies, LED Illumination systems, optics, or cameras^28,36^. Thus, it remains challenging to implement IOSI for labs without specific expertise in microscopy design, optics, or electronics, or those on a budget.

Here, we build on prior IOSI designs to provide a set of simple adaptations that enable the addition of IOSI capabilities to an existing upright microscope. We use an inexpensive, commercially available LED ring light controlled by an Arduino microcontroller for illumination, a basic microscope objective for light collection, and mid-range cameras for detection to achieve high-sensitivity IOSI. In addition, we provide simple, easy-to-use MATLAB based applications for illumination control and basic image analysis. Using this system, we are able to map sensory-evoked signals across primary somatosensory cortex (S1) and primary visual cortex (V1) and we demonstrate that these maps are highly consistent across animals. We also show our IOSI imaging set-up is sufficiently sensitive to detect intrinsic signals from small cortical areas (~200-300 μm in diameter), such as a single barrel in the barrel field (S1BF), and that we can quantify cortical map plasticity longitudinally after whisker trimming. Together, these innovations to the IOSI set up significantly lessen financial and technical barriers for researchers wishing to add this versatile technique in neuroscience to their laboratories.

## Methods

### Experimental Animals

All experiments followed the U.S. National Institutes of Health guidelines for animal research, under an animal use protocol approved by the Chancellor’s Animal Research Committee (ARC) and Office for Animal Research Oversight at the University of California, Los Angeles (#2020-114). Male and female mice were used, beginning at 7-10 weeks old at the time of cranial window surgery. All animals were housed in a vivarium with a 12 h light/dark cycle. For these experiments we used 3 hemizygous transgenic Ai162D mice (Ai162(TIT2L-GC6s-ICL-tTA2)-D, Jax line 031562)^37^, 3 homozygous and 6 heterozygous PV-Cre mice (B6.129P2-Pvalbtm1(cre)Arbr/J, Jax line 017320)^38^, and 1 Thy1-jRGECO1a transgenic mouse (Jax line 030526)^39^. All transgenic lines were maintained on a C57BL/J6 background.

### Cranial Window Surgery

Implantation of chronic glass cranial windows was performed as previously described^40,41^. Mice were deeply anesthetized using 5% isoflurane followed by maintenance with 1.5-2% isoflurane. The scalp was removed and periosteum cleaned away by gentle scraping. An ~4 mm diameter circular craniotomy, centered ~3 mm lateral to the midline and ~2 mm caudal to Bregma was made using a pneumatic dental drill with a FG ¼ drill bit (Midwest Dental) over the primary somatosensory cortex, including the barrel field (S1BF), forelimb (S1FL) and hindlimb (S1HL) areas. For primary visual cortex (V1), the craniotomy location was shifted ~1.5 mm caudal. The craniotomy was sealed using either a single 5 mm #1 sterile glass coverslip (Harvard Apparatus), or a 4 mm coverslip glued to a 5 mm coverslip using an optical adhesive (Norland Products, #71), and was glued to the skull with cyanoacrylate glue (Krazy Glue) followed by dental acrylic (OrthoJet, Lang Dental). A small stainless steel headbar was placed rostral to the cranial window and embedded in dental acrylic to allow subsequent fixation of the mouse onto the microscope stage. Carprofen (5 mg/kg, i.p., Zoetis) and dexamethasone (0.2 mg/kg, i.p., Vet One) were provided for pain relief and mitigation of edema on the day of surgery and daily for the next 48 h. Mice were allowed to recover from the surgery for 3 weeks before the first imaging session.

### IOSI Hardware/Software Setup

We adapted an upright two-photon microscope (Bergamo II, Thorlabs) to perform intrinsic signal imaging. A 4x air immersion objective (Nikon CFI Plan Apochromat Lambda, 0.2 NA) was threaded into the microscope objective holder and a ring of 16 LEDs with integrated drivers (Adafruit NeoPixel Ring, #1463) was affixed to the objective using a custom 3D-printed holder. The ring LED illumination was controlled using an Arduino microcontroller (Uno Rev3) and a custom written MATLAB (Mathworks) application. Full details on how to install and set-up the ring LED illumination, including files for 3D printing and MATLAB code can be found on github (https://github.com/zeigerlab/Intrinsic-Signal-Imaging). Light from the objective was transmitted directly to a camera tube and camera, either a 1x camera tube (Thorlabs WFA4100) coupled to an 8 megapixel CCD camera (Thorlabs, 8051M-USB) or a 0.5x camera tube (Thorlabs WFA4102) coupled to a 12.3 megapixel CMOS camera (Thorlabs, CS126MU). A 5 V TTL pulse generated 1 second prior to the onset of stimuli was used to trigger camera acquisition (Thorlabs, TSI-IOBOB).

### IOSI Acquisition

Animals were sedated with chlorprothixene (~3 mg/kg, i.p.), lightly anesthetized with ~0.5-0.7% isoflurane, and head-fixed below the microscope. The cortical surface was illuminated by green light (525 nm) to visualize and capture an image of the superficial vasculature. The microscope was then focused 300 μm below the cortical surface and red light (625 nm) was used to record intrinsic signals, with frames collected at 10 Hz starting 1 s before and up to 3 s after stimulation onset. Thirty trials with inter-stimulus intervals of 20 s were conducted for each imaging session. Paw and whisker stimuli (10 or 100 Hz sine wave, 1.5 seconds long) were generated in MATLAB, output via a multifunction input/output device (National Instruments, BNC 2090a and PCIe-6363) to a voltage amplifier (Micromechatronics, PD200-V100,100), and delivered using a glass capillary affixed to a piezoelectric bending actuator (Bimitech Python PBA6014-5H200). Visual stimuli were generated using PsychoPy^42^ and consisted of drifting sinusoidal gratings (spatial frequency 0.04 cycles/deg; speed 2 cycles/s) at orientations of 0, 45, 90, and 135° (each displayed for 0.375 seconds for a total stimulus duration of 1.5 seconds) followed by a black screen during the inter-stimulus interval. Images were acquired with 100 ms exposure time at 10 frames per second, for a total of 4 seconds per trial.

### Quantification of Evoked Signals

Evoked signals were quantified as follows. Images were first gaussian filtered and spatially downsampled by a factor of four. For each trial, an average baseline reflectance image was created by calculating the mean across 0.9 s of images (9 frames) prior to stimulus onset. Post-stimulus reflectance images were temporally averaged across 0.3 s bins for 1.5 seconds total, starting 0.5 s after stimulus onset, yielding 5 post-stimulus images per trial. Change in reflectance values (ΔR/R) were then calculated by subtracting the average baseline reflectance from each post-stimulus image and dividing the result by the average baseline reflectance. These ΔR/R values were then averaged across all 30 trials and finally summed across the 5 post-stimulus images to yield a single total stimulus-evoked ΔR/R image. A circular mask corresponding to the area of the cranial window (based on the vasculature image) was fit to the stimulus evoked ΔR/R image such that pixels outside the cranial window were set to 0. Binary images were then created either by thresholding ΔR/R values using a percentage of the maximum signal intensity (for larger maps, such as S1BF or V1), or by first calculating Z-scores of the obtained ΔR/R values and then thresholding values below a Z-score of −3 (for smaller maps, such as single whisker, S1FL, or S1HL). Binarized images were then pseudocolored and overlaid onto images of the vasculature. To obtain group averaged images and map displacements, binarized maps of the S1BF, S1FL, and S1HL were fit with an ellipse using the “Analyze Particles” function in Fiji/ImageJ^43^. The center of each ellipse was determined and stimulus evoked ΔR/R images from each mouse were aligned using the center of the S1BF ellipse, overlaid, and averaged. Displacement of S1FL and S1HL relative to S1BF were then calculated for each mouse and compared using a oneway multivariate analysis of variance (MANOVA). To quantify map area for single whisker evoked maps, a median filter was applied to binarized maps to remove noise and the area of thresholded pixels was calculated. Pre- and post-whisker trimming map sizes were compared using a paired sample, two-tailed t-test. All values listed or plotted are mean ± standard error of the mean, unless otherwise specified.

### Whisker Trimming

Animals were anesthetized with isoflurane (5% for induction, 1.5-2% for maintenance) and all whiskers on the right side of the snout, except those undergoing stimulation (B1,C1, and/or D1), were trimmed using a fine scissors to a length of ~5 mm immediately prior to imaging. For the chronic whisker trimming experiment, all whiskers on the right side of the face except C1 were trimmed flush with the vibrissal pad immediately following baseline imaging and re-trimmed as needed to remove any whisker re-growth, approximately three times weekly, for a total of three weeks.

## Results

To adapt an existing upright microscope for IOSI, we designed a simple, easy to implement system for sample illumination and image acquisition (**Fig. 1A**). The brain surface is illuminated using an array of LEDs and reflected light is collected through a 4x microscope objective and transmitted to a scientific camera without the need for any intervening filters (**Fig. 1B**). For sample illumination, we used a pre-fabricated commercially available LED ring light controlled by an Arduino microcontroller (see Methods). These LEDs have integrated drivers and require only a basic 5 V AC/DC power adapter, obviating the need for custom built circuits or high-end power supplies. The ring light is slipped onto a 4x microscope objective using a simple 3D printed mount and provides even sample illumination at wavelengths of ~470 nm, ~525 nm, or ~625 nm. Control of the illumination wavelength and light intensity is achieved with a MATLAB-based application (**Fig. 1C**). Camera acquisition can then be synchronized to stimuli of interest via a transistor-transistor logic (TTL) pulse from the imaging computer. After image acquisition, images are processed using a flexible MATLAB-based application (**Fig. 1D**) to generate scaled change in reflectance (ΔR/R) images, or further thresholded and overlaid onto an image of the brain vasculature for localization of signals. The image analysis application is flexible with customizable inputs for specific acquisition settings, including imaging framerate, number of trials, baseline and stimulus duration, and temporal binning, among others (**Fig. 1D**). The entire system can be set up on an existing upright microscope in a few hours for a cost of <$5000, or <$50 if the microscope is already equipped with an appropriate objective and camera. Complete installation instructions and design files are freely available online (https://github.com/zeigerlab/Intrinsic-Signal-Imaging).

**Figure 1.**
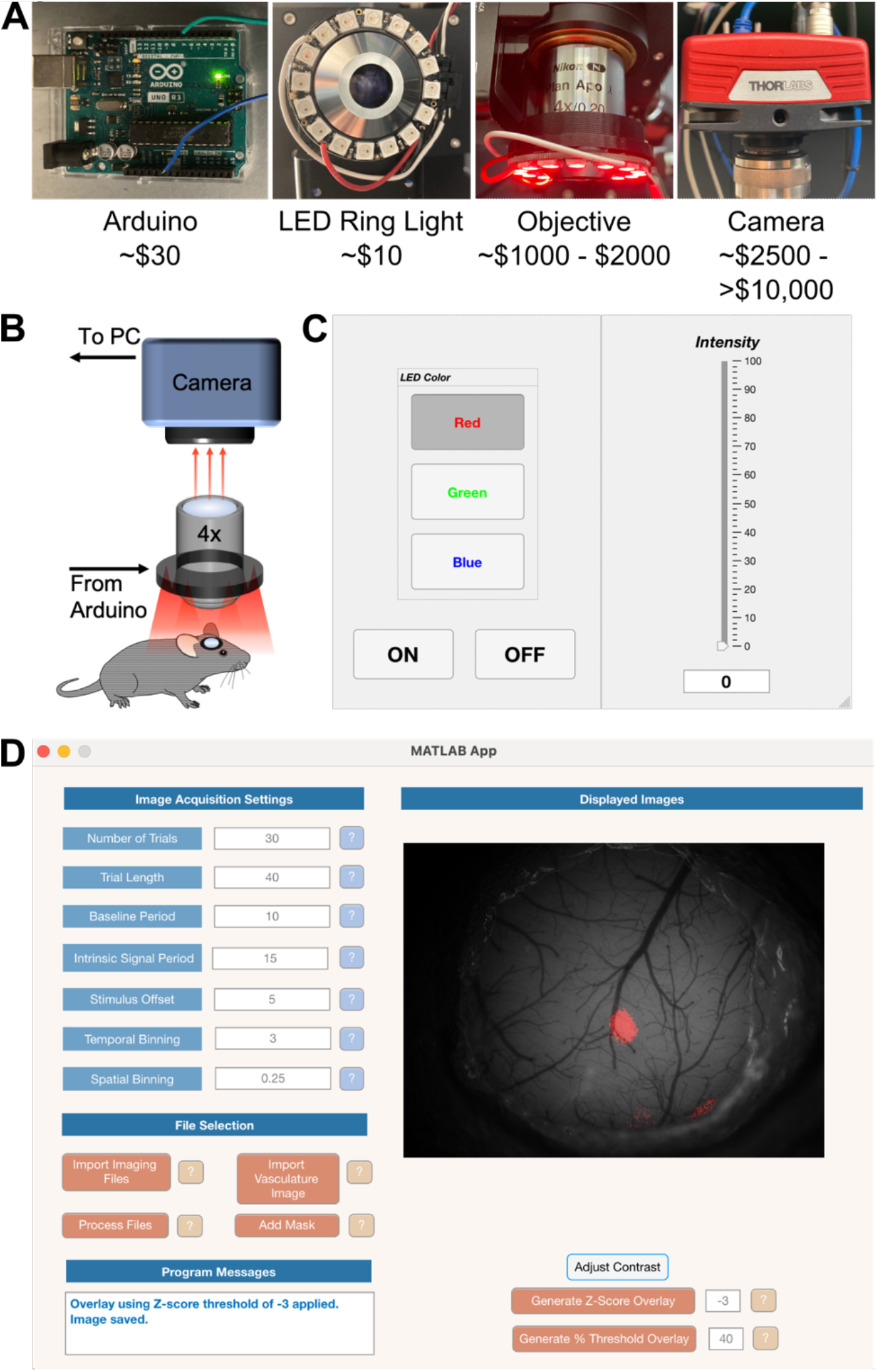
Hardware and Software Adaptations to enable IOSI on an existing upright microscope. **A.** Hardware necessary to achieve IOSI includes (from left to right) an Arduino microcontroller connection to an LED ring light for illumination, an objective for light collection, and a camera to record reflected light. **B.** Schematic of the hardware adaptations and basic connections. **C.** Image of the application for illumination control. Users may set the illumination wavelength and intensity. **D.** Image of the application for basic image analysis. Users may input settings used for image acquisition and identify acquired images. The application will then process images, calculate ΔR/R values and display a scaled image in the application window. Thresholding can then be done within the application, using either an absolute percentage of the maximum ΔR/R value or using Z-scores of ΔR/R values, and a binarized map can then be overlaid onto an image of the vasculature for localization of signals.

IOSI is commonly used to localize areas of the mouse S1, such as the barrel field (S1BF), forelimb (S1FL), or hindlimb (S1HL). Using our simplified set-up for imaging, we performed IOSI through a chronic cranial window implanted over the left primary somatosensory cortex during vibrotactile stimulation of the contralateral whiskers, forelimb, or hindlimb (**Fig. 2A**). We were able to record robustly evoked signals corresponding to the S1BF, S1FL, and S1HL in individual mice (**Fig. 2B**). We then aligned the S1BF signals across 6 different animals and found that the group averaged signal for each sensory area was highly localized (**Fig. 2C**), demonstrating that the location of these sensory-evoked maps is consistent and reproducible across animals. By thresholding and merging the S1BF, S1FL, and S1HL signals, we could clearly define distinct maps that were approximately situated as expected anatomically (**Fig. 2A,D**)^44^. To further define the precision and reproducibility of our obtained maps, we quantified the relative displacement of the center of the S1FL and S1HL maps relative to the center of the S1BF map in each of six mice (**Fig. 2E**). The displacements were tightly clustered by map (S1 FL or S1HL) with the S1FL map displacements significantly distinct from the S1HL map displacements (MANOVA, *p*=1.04 x 10^-5^). We also performed IOSI during presentation of drifting sinusoidal gratings to map the primary visual cortex and again found we were able to identify robust visually evoked signals (**Fig. 2F,G**). Together, these results demonstrate that our IOSI set-up is sufficiently sensitive to record evoked signals from several major sensory areas across the mouse cortex.

**Figure 2.**
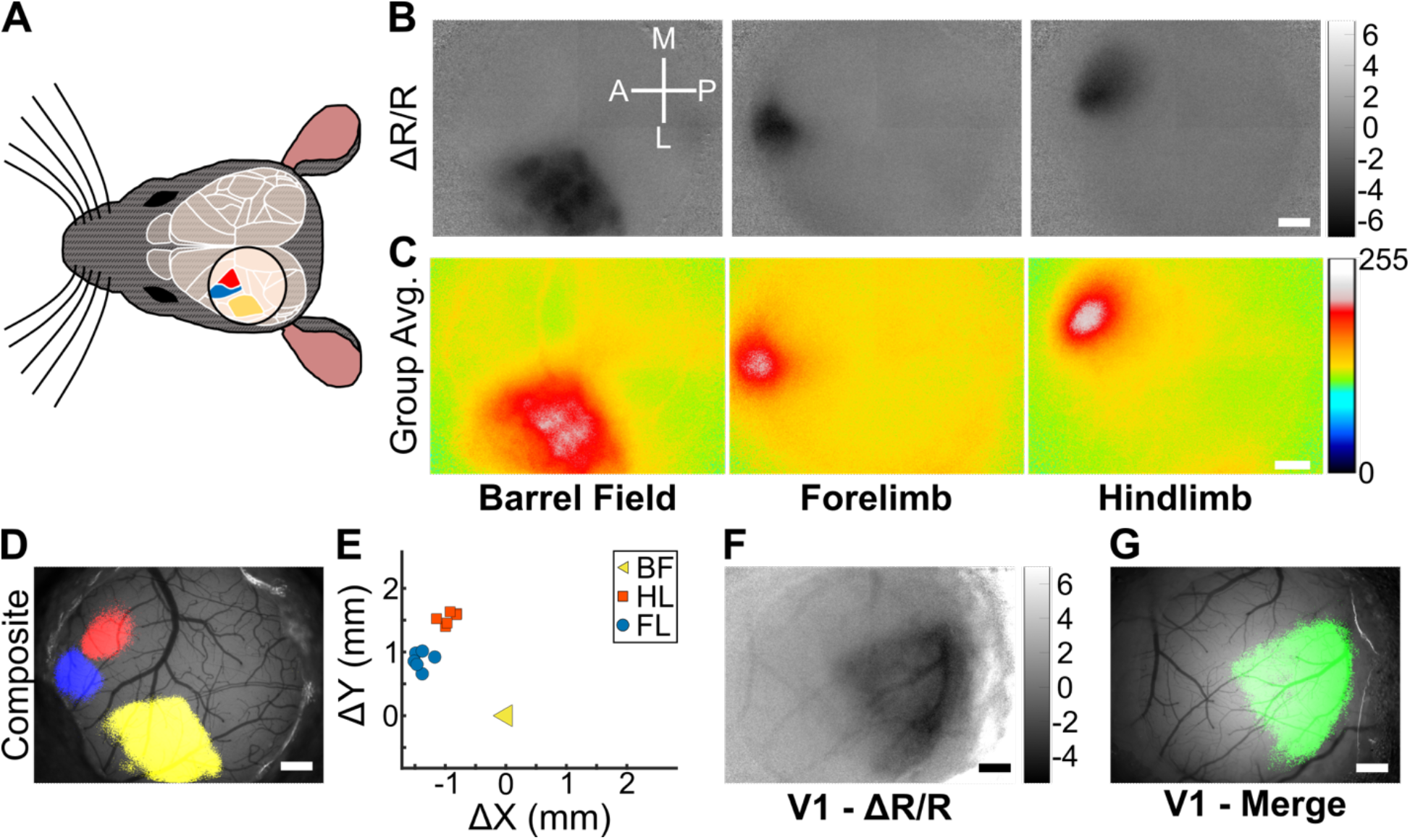
Mapping evoked signals across primary somatosensory and visual cortices. **A.** Schematic of cranial window placement for mapping evoked signals in the primary somatosensory cortex. Approximate locations of major cortical regions are outlined in white (adapted from the Allen Mouse Brain Atlas Brain Explorer 2), with the S1FL (blue), S1HL (red), and S1BF (yellow) color coded. **B.** Scaled ΔR/R images of IOSI from an individual mouse performed during vibrotactile stimulation of the contralateral whiskers, forelimb, or hindlimb. Scaled ΔR/R values are x 10^-4^. Scale bar = 0.5 mm. **C.** Group averaged images from 6 mice of IOSI performed during vibrotactile stimulation of the contralateral whiskers, forelimb, or hindlimb. Values are arbitrary units of 8 bit images from minimum (0) to maximum (255). Scale bar = 0.5 mm. **D.** Merged sensory-evoked maps from (B) overlaid onto the cortical vasculature. ΔR/R images were Z-scored and thresholded for values <-3, binarized, pseudocolored (S1FL [blue], S1HL [red], and S1BF [yellow]), then merged and overlaid. Scale bar = 0.5 mm. **E.** Displacement of S1FL and S1HL map centers, relative to the S1BF map, from 6 individual mice. Map displacements for S1FL and S1HL were significantly different from one another (one-way MANOVA, p=1.04 × 10^-5^). **F.** Scaled ΔR/R images of IOSI from an individual mouse performed during passive viewing of drifting sinuosodal gratings by the contralateral eye. Scaled ΔR/R values are x 10^-3^. Scale bar = 0.5 mm. **G.** Visual-evoked map from (F) overlaid onto the cortical vasculature. Scale bar = 0.5 mm.

We next sought to determine if we could record evoked signals from smaller cortical areas. The mouse S1BF exhibits strong somatotopic organization, with sensory signals from individual whiskers predominantly encoded within single cortical columns, referred to as barrels, that are ~200-300 μm in diameter^45^. We performed IOSI during vibrotactile stimulation of single whiskers (either the B1, C1, or D1 whisker) and found we were able to record robust single-whisker evoked signals (**Fig. 3A,B**). After thresholding and binarizing to obtain single-whisker maps, we merged the three individual single-whisker maps and found these were clearly distinct and adjacently arranged according to the expected somatotopic organization of the S1BF (**Fig. 3C**). To quantify the minimum number of trials necessary to obtain reliable single-whisker maps, we calculated the cumulative change in reflectance values (ΔR/R) across each of the 30 trials performed in an individual IOSI session for 18 single-whisker maps (3 whiskers in each of 6 mice). We then calculated the area of the evoked signal for each trial as a percentage of the maximum area obtained using all 30 trials (**Fig. 3D**). We found that map area plateaued after 25-30 trials, but was already greater than 50% of the total map area after as few as 10 trials. Finally, we also confirmed that we could generate similar single-whisker maps to those obtained with a CCD camera using a more cost-efficient CMOS camera (**Fig. 4**). Thus, our IOSI set-up is sufficient to obtain reliable sensory-evoked maps with high sensitivity even from signals generated by relatively small areas of cortex.

**Figure 3.**
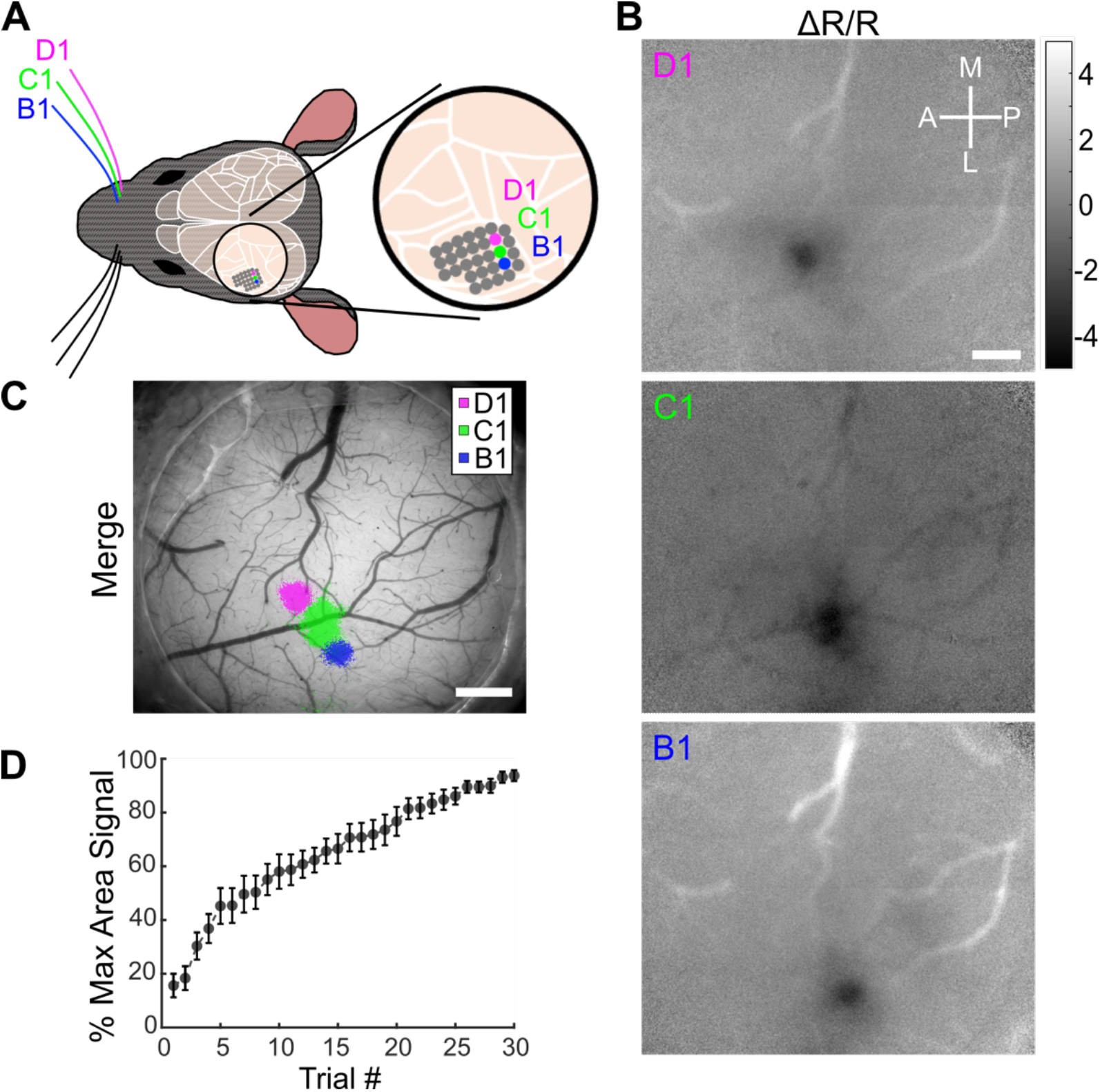
IOSI of single whisker evoked cortical activity in the S1BF. **A.** Schematic of stimulated whiskers (B1, C1, and D1) and somatotopic organization of the corresponding barrels in the S1BF. **B.** Scaled ΔR/R images of IOSI from an individual mouse performed during vibrotactile stimulation of the contralateral B1, C1, and D1 whiskers. Scaled ΔR/R values are x 10^-3^. Scale bar = 0.5 mm. **C.** Merged single whisker maps from (B) overlaid onto the cortical vasculature. ΔR/R images were Z-scored and thresholded for values <-3, binarized, pseudocolored, then merged and overlaid. Scale bar = 0.5 mm. **D.** Average map area by IOSI trial, as a percentage of the maximum map area following all 30 trials, of 18 single whisker representations in 6 mice.

**Figure 4.**
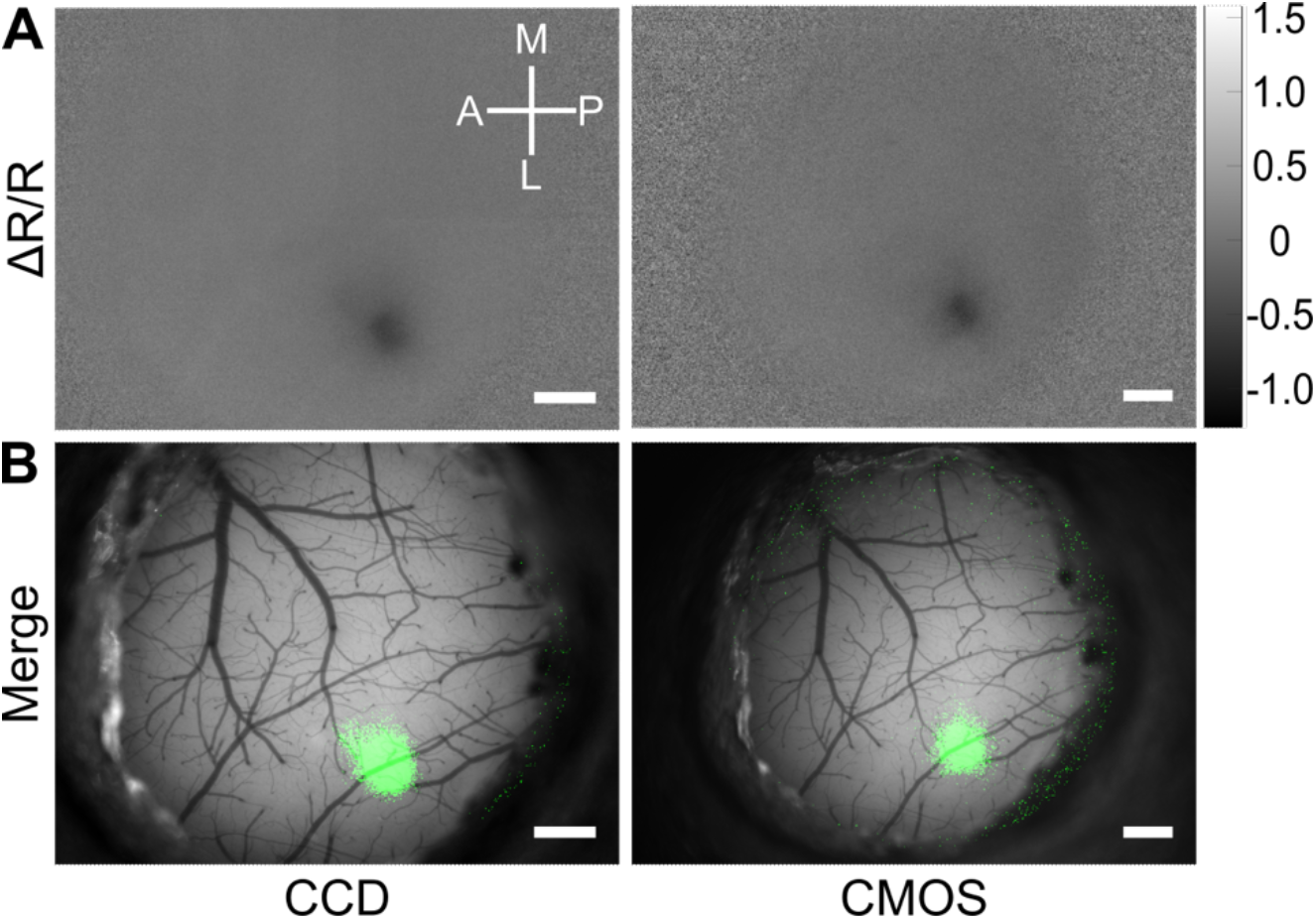
High sensitivity IOSI can be achieved with CCD or CMOS cameras. **A.** Scaled ΔR/R images of IOSI from an individual mouse performed during vibrotactile stimulation of the contralateral C1 whisker. Scaled ΔR/R values are x 10^-2^. Scale bar = 0.5 mm. **B.** Merged single whisker maps from (A) overlaid onto the cortical vasculature. ΔR/R images were Z-scored and thresholded for values <-3, binarized, pseudocolored, then merged and overlaid. Scale bar = 0.5 mm.

In addition to merely localizing signals, IOSI has also frequently been used to study the plasticity of cortical maps over time. One of the most commonly used paradigms for inducing cortical map plasticity in rodents is whisker trimming. Following chronic trimming of whiskers, cortical map areas corresponding to the spared whisker expand, whereas those corresponding to trimmed whiskers shrink^14,17,46^. Having shown that our IOSI setup is sufficiently sensitive to measure single-whisker maps, we tested whether we could use our set-up to measure plasticity of these single whisker maps in response to whisker trimming. Following baseline IOSI imaging of C1 whisker evoked maps (**Fig. 5A,B** left panels), we chronically trimmed all whiskers of the contralateral vibrissal pad except C1. Three weeks later we repeated IOSI during C1 whisker stimulation and quantified the area of the evoked C1 map (**Fig. 5A,B** right panels). We merged the resulting pre- and post-whisker trimming maps and found that the C1 whisker map location was stable, but had increased in size, as expected, following chronic whisker trimming sparing only the C1 whisker (**Fig. 5C**). To quantify this effect, we calculated the C1 whisker evoked map area for 6 mice pre- and post-trimming and found an ~82% increase in map area after trimming (**Fig. 5D,** pre-trimming = 0.16 ± 0.03 mm^2^, post-trimming = 0.30 ± 0.03 mm^2^, paired sample, two tailed t-test *p*=0.003). Thus, our IOSI set-up yields highly sensitive, quantitative cortical maps that can be used to both localize cortical signals as well as study fundamental processes such as cortical map plasticity.

**Figure 5.**
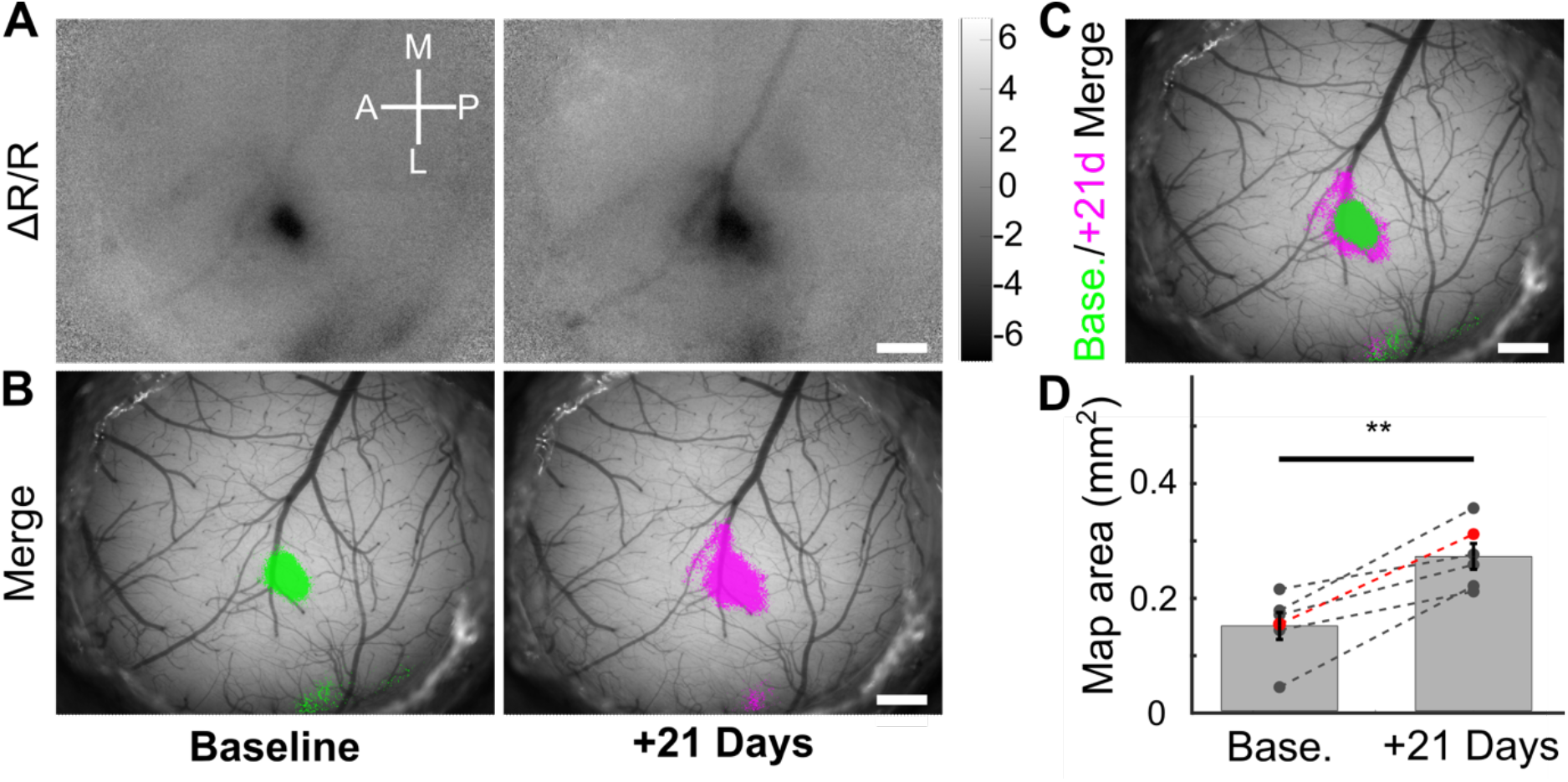
Longitudinal IOSI detects cortical map plasticity following whisker trimming. **A.** Scaled ΔR/R images of IOSI from an individual mouse performed during vibrotactile stimulation of the contralateral C1 whisker at baseline before (left) and 21 days after whisker trimming sparing only the C1 whisker. Scaled ΔR/R values are x 10^-3^. Scale bar = 0.5 mm. **B.** Merged single whisker maps from (A) overlaid onto the cortical vasculature. ΔR/R images were Z-scored and thresholded for values <-3, binarized, pseudocolored, then merged and overlaid. Scale bar = 0.5 mm. **C.** Merged pre- and post-whisker trimming maps, aligned using vasculature images, showing expansion of the C1 whisker cortical map representation. Scale bar = 0.5 mm. **D.** Quantification of change in C1 map area after whisker trimming from 6 mice. The dots in red represent the mouse depicted in panels A-C. **, paired sample, two-tailed t-test, *p*=0.003.

## Discussion

Here we describe a set of simple adaptations that can be applied to an existing upright microscope to allow high sensitivity IOSI. These adaptations make use of readily available commercial parts, can be implemented in an afternoon (or less), require no custom machining, and do not require any specialized skills in optics, electronics, circuit design or programming. In addition, the entire set-up can be implemented at extremely low cost - ~$5,000 for an entire set-up or <$50 if the microscope to be adapted is already equipped with a sufficient camera and objective. In addition, we provide simple MATLAB-based applications for controlling illumination and processing acquired images to obtain raw change in reflectance signals or localized maps overlaid onto images of the cortical vasculature. Using this set-up we were able to record strong evoked signals from the major areas of the mouse somatosensory cortex, including the S1BF, S1FL, S1HL, and V1. These maps show excellent signal-to-noise, can be clearly distinguished from adjacent cortical areas, and are highly reproducible across mice. We were also able to record sensory-evoked signals from smaller cortical areas, at least as small as a single cortical barrel. Indeed, we could consistently generate clear single-whisker evoked cortical maps from several adjacent whiskers that aligned somatotopically as expected in the cortex. Furthermore, we found that our IOSI set-up could be used to quantitatively measure changes in cortical map plasticity. By tracking single-whisker evoked maps over time following whisker trimming, we found a significant increase in the cortical map area of the spared C1 whisker.

The adaptations we describe here update and simplify existing methods for IOSI in several ways. At the most basic level, a system for performing IOSI requires a light source for illuminating the cortical surface, optics to collect reflected light, and a camera to record the collected light^1,2^. Early IOSI imaging set-ups utilized traditional broadband light sources integrated into the micro- or macroscope housing, collimated and passed through excitation filters to achieve illumination^4,34^. Later iterations incorporated external light sources on flexible light guides to simplify the construction of a micro/macroscope, but this requires subjective adjustments on a sample-by-sample basis by the user to achieve uniform sample illumination^28,35^, potentially reducing reproducibility. More recently, arrays of LEDs coupled to the collection optics have been used, but these were often custom built, requiring machining, custom circuit fabrication, and high-end power supplies to achieve stable illumination^36^. The LED ring light we describe here holds several advantages: 1) it is commercially available at low cost; 2) it allows for three color illumination (including 525 nm for visualizing vasculature and 625 nm for IOSI); 3) it has integrated drivers and can be controlled with a basic Arduino microcontroller; 4) it is powered by a simple 5V AC/DC power adapter plugged directly into a standard electrical outlet; and 5) it can be easily attached to the microscope using a simple 3D printed mount. Further, we have created an easy-to-use MATLAB-based application to control the illumination. Together, these features allow for stable, even illumination of samples that can be connected with just a few wires, with no specialized skills in optics, electronics, or programming required.

For collection optics, we recommend an off-the-shelf, commercially available 4x plan apochromatic microscope objective. Traditionally, a tandem lens macroscope is often incorporated into IOSI set-ups for light collection, as the tandem lens provides an excellent combination of field-of-view (FOV), working distance, and numerical aperture^34^. However, as with illumination technology, microscope optics have advanced since the tandem-lens configuration was first introduced, and now even relatively inexpensive microscope objectives have numerical apertures sufficient to achieve IOSI (NA of ~0.1-0.2, compared to ~0.4 for a tandem lens configuration). We found that qualitatively, single whisker maps obtained using this set-up with a 4x microscope objective were at least as robust as those these authors have obtained in the past using a tandem lens macroscope configuration^32^. For cameras, we tested both a CCD and a CMOS scientific microscope camera. Since intrinsic signals are small, ~0.01-0.1% of total reflected light, having a camera with sufficient sensitivity is essential, with bit depth of 10 or greater, low noise, and high full well capacity often cited as some of the most important characteristics^35^. The cameras we tested here had specifications (and costs) in the mid-range of available scientific cameras in the same class (CCD vs CMOS) and both were able to achieve qualitatively similar results mapping single-whisker evoked responses.

While our IOSI set-up has many advantages, there are some limitations. The LED ring light we have used here will not fit on tandem lens macroscopes or significantly larger microscope objectives. However, the same LEDs are available in a variety of additional forms, including rings with larger diameters and strings of LEDs that can be shaped into custom forms. As such, it should be possible to adapt the same illumination strategy, including the same wiring, microcontroller, and illumination control application, across set-ups with varying configurations of collection optics. Our illumination control application works well for constant single-wavelength illumination, but is not set-up for multi-spectral imaging^47,48^. It should be possible to achieve the rapid wavelength switching synchronized to camera acquisition using the same hardware we have described, but this will require additional customization on the part of individual users. To date, we have used two cameras in our experiments, but we have not exhaustively confirmed that IOSI can be achieved equally well with all available scientific cameras. We believe a wide range of CCD and CMOS cameras will likely function interchangeably given that the tested cameras have relatively modest midrange technical specifications, but refer readers to additional resources for factors to consider when choosing a particular camera for their imaging set up^1,36^. Regarding field-of-view, with a 4x objective, we achieved a FOV of ~4.5 x 3.4 mm with the CCD camera, and ~5.7 x 4.3 mm with the CMOS camera used in this study. These FOV sizes are sufficient to visualize the most commonly used craniotomy sizes in mice, but may not be sufficient for bilateral cortex wide imaging or imaging larger animals. However, by reducing the magnification on the microscope objective or camera tube lens, utilizing a larger format camera sensor, and/or switching to a tandem lens macroscope configuration, it is possible to achieve a significantly larger FOV. Our imaging processing application has been designed with flexibility in mind, allowing users to adjust many settings that might be specific to their particular imaging set-up (e.g. framerate, duration of imaging, temporal binning, intrinsic signal of interest). However, we cannot guarantee functionality for more custom use-cases. In addition, the application generates scaled ΔR/R images and maps overlaid onto vasculature, but users may require more advanced image processing tools in certain cases (e.g. retinotopic mapping of higher visual cortical areas). Finally, we have not provided detailed instructions here on generating stimuli to evoke cortical signals as we anticipate these will vary significantly from laboratory to laboratory based on individual experimental paradigms. We have example code and set-up instructions for the piezoelectric vibrotactile and visual stimuli used in our laboratory available online (https://github.com/zeigerlab/Intrinsic-Signal-Imaging).

We envision these adaptations will be best suited to laboratories looking to add IOSI capabilities to an existing upright microscope in the laboratory. The primary advantages include low cost, ease of implementation, and small footprint given that a separate dedicated IOSI rig is not required. However, we anticipate these updates may be useful for more advanced users as well who might want to update components of existing rigs or incorporate these adaptations into custom-built stand-alone IOSI rigs.

## Acknowledgements

We thank Thorlabs for loaning the CMOS camera used in this study and Dr. Carlos Portera-Cailliau for advice and helpful feedback. This work was supported by National Institutes of Health Grant 1K08NS114165-01A1 and American Academy of Neurology Grant NRTS 2199.

